# Adolescent female rats undergo full systems consolidation of an aversive memory, while males of the same age fail to discriminate contexts

**DOI:** 10.1101/2020.08.14.250860

**Authors:** Ana P. Crestani, Fernanda N. Lotz, Mirelle A. Casagrande, Bruno Popik, Kétlyn T. K. Guerra, Lucas de Oliveira Alvares, Jorge A. Quillfeldt

## Abstract

Generalization is an adaptive process that allows animals to deal with threatening circumstances similar to prior experiences. Systems consolidation is a time-dependent process in which memory loses it precision concomitantly with reorganizational changes in the brain structures that support memory retrieval. In this, memory becomes progressively independent from the hippocampus and more reliant on cortical structures. Generalization, however, may take place much faster in adult animals depending on the presence of sex hormones. Notwithstanding its relevance, there are few studies on sex differences in memory modulation. Here, a contextual fear discrimination task was used to investigate the onset of memory generalization and hippocampus-independence in adolescent male and female rats (P42-49). Subjects were tested 2, 7, 14, 21 or 28 days after training, with females showing memory generalization from day 21 on, whereas males surprisingly unable to discriminate contexts at any time. Ovariectomized females, however, displayed an early onset of generalization. Consistently, pre-test pharmacological blocking of dorsal hippocampus was able to impair memory retrieval in females, but not in males, which indicate that precise memory is dependent on the hippocampus. To our notice, this is the first report of a memory systems consolidation process – expressed in its two dimensions, neuroanatomical and qualitative – in adolescent female rats, and one that can also be accelerated by the reduction of sex hormones through ovariectomy. It is also unprecedented that despite adolescent male rats being able to remember fear learning, they did not discriminate contexts with any precision.

## INTRODUCTION

The ability to make associations between places and events emerges already in infancy (J W Rudy & Morledge, 1994) and predicting the safety of future experiences provides adaptive advantages – a capacity that might also have had evolutionary consequences (Jasnow et al., 2017). Several important modifications occurring in the nervous and endocrine systems during ontogeny may have considerable influence on how detailed memories can be produced, and to what extent animals can discriminate between similar, yet different environments (Spear, 2000).

Previous studies have demonstrated that memory loses precision in a contextual discrimination task as the interval between training and test increases: as time after training passes, it becomes more generalized (de Oliveira Alvares et al., 2012; Wiltgen et al., 2010; Wiltgen and Silva, 2007). Modifications in brain regions that support memory retrieval also occur over time, with retrieval of the memory trace becoming progressively independent from the hippocampus and reliant on cortical areas (Bontempi et al., 1999; Crestani et al., 2018; Frankland et al., 2004; Martin et al., 2005; Maviel et al., 2004; Quillfeldt, 2019; Quillfeldt et al., 1996). Despite not fully understood, systems consolidation appears to express itself in two different dimensions, one neuroanatomical – the gradual independence from the hippocampus – and the other, qualitative – the progressive inability to discriminate between similar contexts, i.e., display memory generalization (Cullen et al., 2015; de Oliveira Alvares et al., 2012; Rudy, 2009; Wiltgen et al., 2010).

Most experiments evaluating memory generalization and systems consolidation are typically performed in adult male rodents, which usually develop over a time-course of around one month (de Oliveira Alvares et al., 2012; Haubrich et al., 2016a; Wiltgen et al., 2010). However, memory generalization can also be experimentally induced in acute form, at any time, though different manipulations such as, e.g., sex hormone levels (Jasnow et al., 2017), stress (Dos Santos Corrêa et al., 2021) or GABA-B receptor manipulation (Lynch et al., 2017). In female rats, for instance, memory generalization was shown to be induced by estradiol. Gonadal hormones are produced in a sex-specific manner throughout adolescence (Cyrenne & Brown, 2011; McCormick & Mathews, 2010; Parker & Mahesh, 1976; Tolson & Chappell, 2012) and clearly influence behavioral performance (Andersen, 2003a; Brown et al., 2015; Doremus-Fitzwater et al., 2012; Spear, 2000; Vetter-O’Hagen & Spear, 2012). Accordingly, brain structures critically involved in learning and memory processes such as the hippocampus, amygdala, and medial prefrontal cortex, undergo extensive morphological and functional remodeling during this period (Andersen, 2003b; Casey et al., 2008), which in the case of context learning might even involve dorsal and ventral hippocampus in different ways (Barfield et al., 2017; Heroux et al., 2019).

Considering the adaptive importance of generalization in the face of potentially threatening situations and the well-known differences in risk-taking behavior between adolescent male and female mammals (Daughters et al., 2013; Doremus-Fitzwater et al., 2012), it is reasonable to suppose differences could also be seen in the time-course of memory generalization observed along the systems consolidation process. In this work, we investigated contextual fear discrimination in adolescent rats (P42-P49) of distinct sexes at different training-test intervals (2d, 7d, 14d, 21d or 28d). Pharmacological inactivation of the hippocampus during retrieval was used to verify hippocampus dependency. Additionally, the possible involvement of female hormones producing the observed effects was confirmed by comparison of intact with ovariectomized females.

## MATERIAL AND METHODS

### Animals

Subjects were the offspring of Wistar rats obtained from the University’s breeding colony. Male and female pups were bred in our housing facility, weaned on postnatal day 21 (P21) and housed with same-sex littermates up to 5 per cage, until their late adolescence and throughout all experimental procedures: between days P42 and P49, these late adolescent rats were trained in a contextual fear conditioning task. Pretraining handling was considered unnecessary since they were familiarized with our facility and the experimenters from the very beginning. Animals were kept on a 12/12h dark/light cycle and had food and water available *ad libitum*. Experiments were conducted during the light cycle (8 a.m. – 6 p.m.) and in accordance with National Animal Care Legislation and Guidelines (Brazilian Law 11794/2008) and approved by the University’s Ethics Committee (project # 23957 and 27143).

### Behavioral Procedure

*Contextual Fear Conditioning* (CFC). CFC chamber consisted of an illuminated Plexiglas box (25.0 × 25.0 cm with a grid floor consisting of parallel 0.1 cm caliber stainless steel bars spaced 1.0 cm apart) with fan background noise. In the conditioning session, rats were placed in the chamber for 3 min, after which they received two 2-sec 0.7 mA footshocks separated by a 30-sec interval. Animals were kept in the conditioning environment for an additional 30 seconds before being returned to their homecages. Three experiments were performed: (i) time-course of the onset of memory generalization in adolescent males and females, (ii) generalization in ovariectomized females compared to intact females and males, and (iii) hippocampal dependency for discriminatory memory retrieval in adolescent animals of both sexes, as shown by local muscimol infusion. Each time point in experiment 1 is a different experimental group, thus each group was trained and tested just once. Test was performed with the animal exposed either to the very same training context, or to a novel one, during 4 minutes in the absence of the unconditioned stimulus (shock). The novel context was a rectangular box 2/3 of the size of the conditioning context, with a smooth floor, without background fan noise and dimmer over-head lighting – also, the chamber was located in a different room, with different spatial cues. For a comparison between contexts A (training) and B (novel) see **Figure 1.N**.

**Figure 1.**
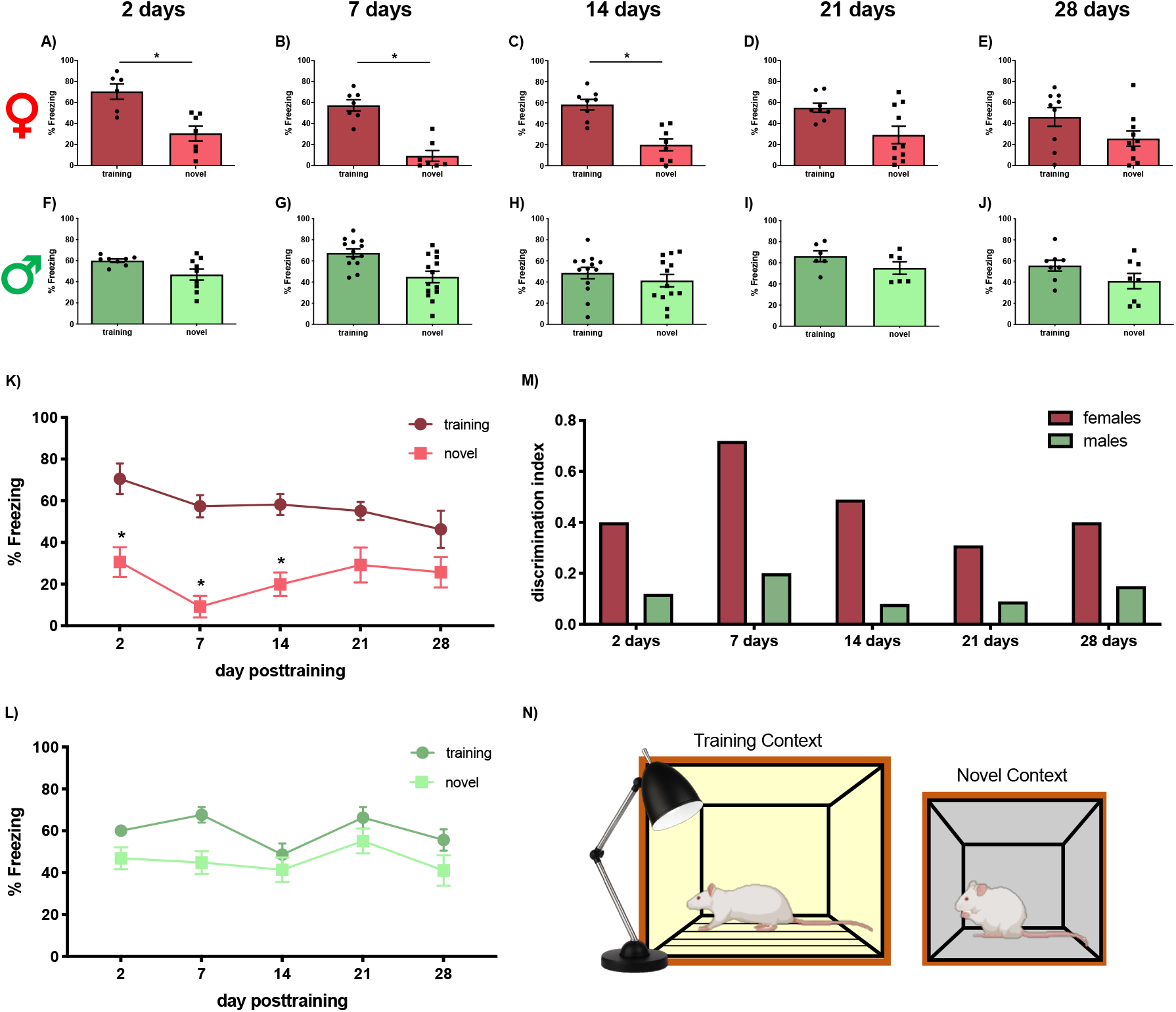
Time-course of fear memory generalization in adolescent female and male rats. Different groups of adolescent female rats were able to discriminate between training or the novel context when tested at **(A)** 2 days (N = 6,7), **(B)** 7 days (N = 7,7) and **(C)** 14 days (N = 8,8) after training. However, memory generalization was verified when tested **(D)** on day 21 (N = 8,10) and **(E)** 28 (N = 9,10). Male rats were tested at the same time points and presented generalization at **(F)** 2 days (N = 8,9), **(G)** 7 days (N = 13,14), **(H)** 14 days (N = 13,13), **(I)** 21 days (N = 6,6) or **(J)** 28 days (N = 8,8) after training. Color code: training context is represented by dark red (females) and dark green (males) while novel context is represented by light red (females) and light green (males). For better visualization, the same results of above bar graphs (A-J) are displayed as line graphs over time, respectively, **(K)** for females, and **(L)** for males. **(M)** Histogram showing a mean discrimination index for males and females in all the time points: index is defined as (% freezing in the training context minus the % freezing in the novel context) / (% freezing in the training context + % freezing in the novel context). Percentage of freezing time was expressed as Mean ± SEM (**A-L**). **(N)** Schematic representation of training versus novel contexts used in the discriminatory tests. (*) P ≤0.05 (see text for more details).

#### First set of experiments: Time-course of contextual fear memory generalization in adolescent males and females

In order to investigate fear memory expression in different, yet similar contexts, adolescent male and intact, naturally cycling female rats were tested in the training or the novel context on days 2, 7, 14, 21 or 28 after training (distinct groups of animals were used at each time point tested). Freezing behavior was recorded in real-time during the 4-min test session by an experienced observer, and was defined as the absence of all movements except those related to breathing. A discrimination index (DI) was calculated aiming to highlight which group displayed the best performance in the test sessions, with DI defined as: DI = (% freezing in the training context minus the % freezing in the novel context) / (% freezing in the training context + % freezing in the novel context). No female animals were monitored for the estrous cycle stage since (a) the procedure was intrinsically stressful and affected performance, and (b) there was no equivalent procedure for males that allowed for the systematization of this behavioral impact; actually, since it is hard to pinpoint the onset of the first cycle with exactitude, females could be in any estrous stage at any of the studied time points, and the homogeneity of data dispersion observed seems to corroborate this.

#### Second set of experiments: Dependence of retrieved memory precision on sex hormone levels 14 days after training

A group of previously ovariectomized (OVX) females was trained in the CFC and tested 14 days after, either in the training or the novel context, in order to evaluate discrimination memory compared to males and intact, naturally cycling females. The time point chosen for the test was 14 days after training since this was the last moment intact females displayed good context discrimination memory while in males generalization was already present, and aimed to evaluate whether precision demanded a sustained level of female sex hormones. The intact female group was basically the same shown in Figure 1.H, thus estrous cycle was not monitored (for the reasons explained above).

##### Ovariectomy procedure

prepubescent rats (P30-33) were anesthetized with an IP combination of ketamine/xylazine (75 and 10 mg/Kg, respectively). A ventral midline skin incision approximately 1.5 cm long was made with subsequent removal of the ovaries, after which the uterine tubes were ligated. Afterwards, ventral muscles and skin were sutured and animals received post-surgery treatment with 200 mg/ml of the analgesic acetaminophen (brand name Tylenol®) diluted in water for 3 days.

#### Third set of experiments: Dependence of retrieved memory precision on hippocampus integrity in males and females 14 days after training

Male and female animals were submitted to stereotaxic surgery to allow for the pharmacological inactivation of the CA1 area of the hippocampus (bilaterally) during contextual discriminatory memory retrieval. The time point chosen for hippocampal inactivation was 14 days after training since this was the last moment intact females displayed good context discrimination memory while in males generalization was already present, and aimed to evaluate whether precision demanded the integrity of the hippocampus. Both males and females were cannulated between the training and the test session (see details below), an invasive surgical procedure that explains the generally lower test % freezing levels observed in the control groups. Also here, to avoid any further stress, the estrous cycle was not monitored in the intact female group, but notice the similar performance levels of vehicle-injected animals of both sexes. In the test session, the experimenter was kept blind to the treatment received.

##### Stereotaxic surgery and cannulae placement

One week after training, rats were anesthetized with an intraperitoneal injection of ketamine/xylazine (75 and 10 mg/Kg, respectively) and submitted to stereotaxic surgery to bilaterally implant guiding cannulae (22-gauge) aimed at the dorsal hippocampus in the following coordinates: AP -3.0 mm (from Bregma), LL ± 2.0 mm, DV -1.5 mm (males) and DV -1.4 mm (females) - DV, i.e., positioned 1.0 mm above target point in each brain structure (Paxinos & Watson, 2007). After surgery, animals were given 6-7 days to recover before being submitted to the behavioral testing procedure, which took place 14 days after training.

##### Drug

Muscimol (1µg/µl; Research Biochemicals International), a GABA_A_ agonist, was dissolved in phosphate-buffered saline (PBS) and bilaterally infused into dorsal hippocampus 15min before test. Concentration (1µg/µl) was chosen to be the exact same employed in previous works with adult males (de Oliveira Alvares et al., 2012, 2013). Vehicle groups went through the same manipulation but received PBS.

##### Intracerebral infusion

At the time of infusion, a 27-gauge infusion needle was inserted into each guide cannula, with its tip protruding 1.0 mm beyond the tip of the cannula and aimed at the dorsal hippocampus. A volume of 0.5 µl was bilaterally infused at a slow rate (20 µl/h), and the needle was removed 30 s after the end of the infusion.

##### Histology

Cannula position was verified for every subject in all experiments. Brains were dissected, immersed in a fixation solution of 20% sucrose and 4% paraformaldehyde, and after a few days were sliced in a cryostat (50µm coronal sections). Sections were stained with cresyl violet and subsequently examined to verify the location of the cannulae. Only animals with correct cannulae placements were considered in the statistical analysis. **Figure 3B-C** shows the typical cannulae placements, histologically verified in the dissected brains.

**Figure 2.**
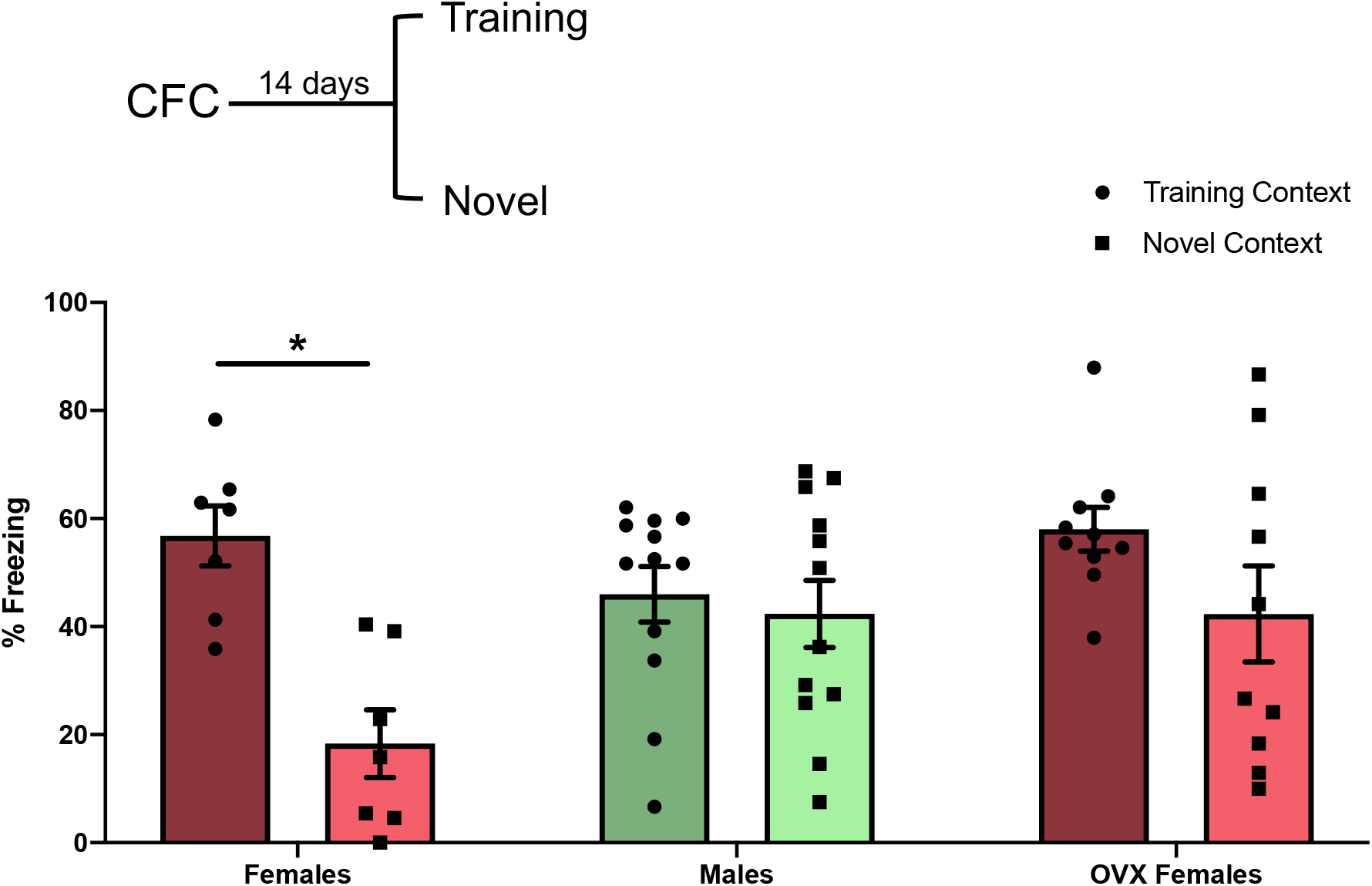
Ovariectomy anticipates memory generalization in female rats. Ovariectomized females (training context N = 10, novel context N = 10) generalize memory earlier then intact females (training context N = 7, novel context N = 7) and males (training context N = 12, novel context N=12). Groups were compared using two-way ANOVA and Tukey’s post-hoc test was used to indicate differences between groups. Percentage of freezing time was expressed as Mean ± SEM. (*) P ≤0.05 (see text for more details).

**Figure 3.**
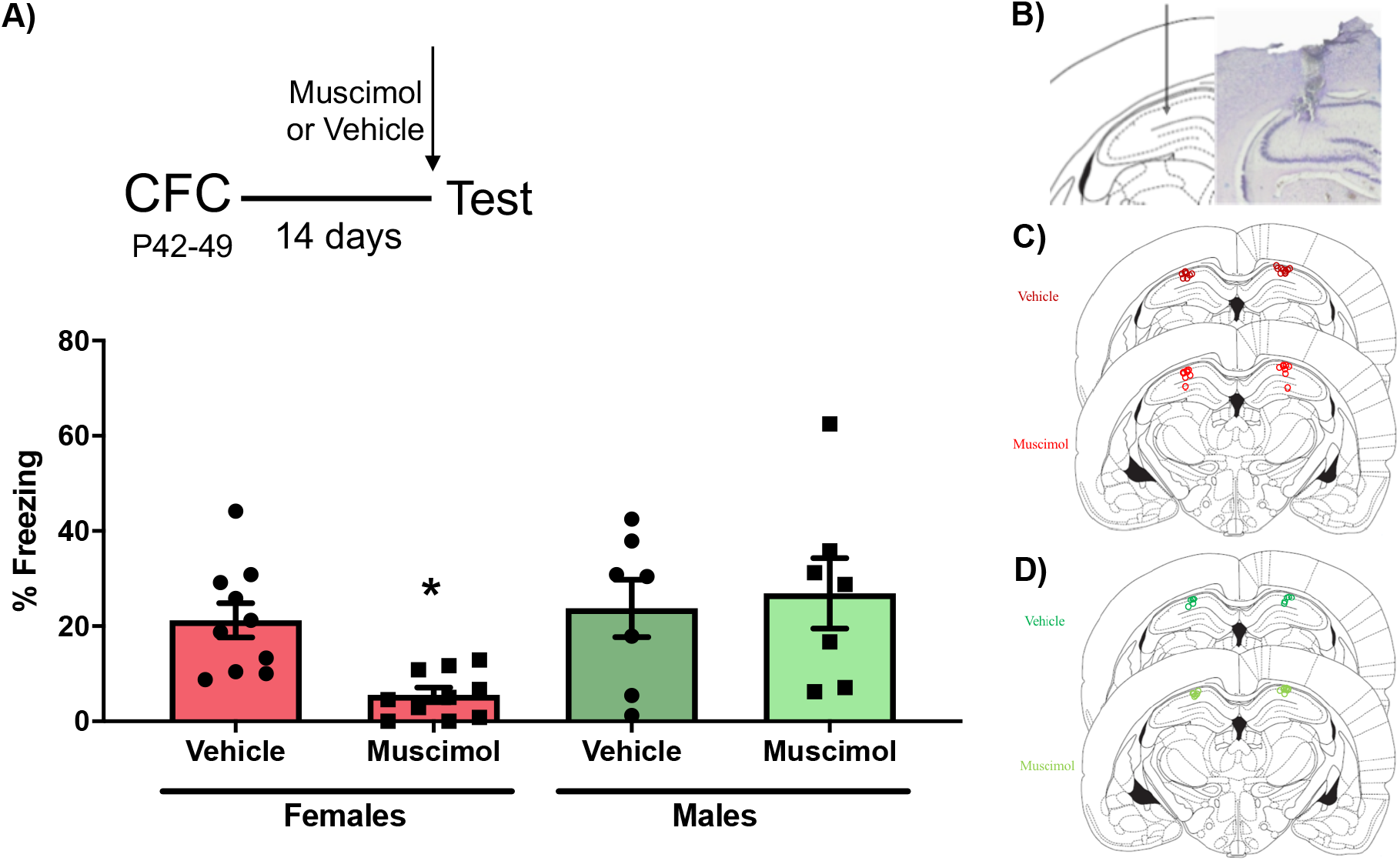
Memory retrieval 14 days after training is dependent on hippocampus in females, but not in males. **(A)** GABA_A_ agonist muscimol was infused in the dorsal hippocampus prior to the retrieval test in the training context. Muscimol impaired memory retrieval in females (N = 10,10), but did not inhibit memory retrieval in males (N = 7,7). Groups were compared using two-way ANOVA followed by Tukey’s post-hoc test. Percentage of freezing time was expressed as Mean ± SEM. (*) Group differs from all other groups with P ≤ 0.05 (see text for more details). **(B)** Confirmation of the positions of the canullae tips of subjects showing a **(B)** histological image with a typical cannula lesion and position; **(C)** cannulae placement in female subject; **(C)** the same for male subjects.

### Statistical analysis

To support the use of parametric statistics, both normality (Kolmogorov-Smirnov Test) and homoscedasticity (Levene test) were checked. In the first experiment, three variables (sex, time and context) were compared by a three-way ANOVA, and the second and third experiments were analyzed by two-way ANOVA, looking for interactions and complemented by main effects for one or more factors. We performed additional comparisons using Tukey HSD test in order to identify those specific experimental groups in which the contexts (experiments 1 and 2) or treatments (experiment 3) actually differ significantly. Although such a procedure is debatable, our intention was only to highlight some specific findings, supporting them with their Ps, and – where appropriate – Cohen’s d effect sizes.

### Transparency and Openness

The sample size was determined based on similar previous studies from our lab (Quillfeldt et al., 1996; de Oliveira Alvares et al., 2012). No animals were excluded from analyses. All data (expressed in percentage of freezing during test session) and statistics are available at Figshare [https://doi.org/10.6084/m9.figshare.14903325.v1]. Data were analyzed using Prism, version 8.

## RESULTS

### 3.1. Time-course of contextual fear memory generalization in adolescent females and males

To test whether sex differences during adolescence influence contextual discrimination, adolescent female or male rats were trained in the contextual fear conditioning and separate groups were tested in either the conditioned, or the novel environment 2, 7, 14, 21, or 28 days later.

Three-way ANOVA showed sex-context interaction (F_(1,158)_ = 13.55, P = 0.0003) and main effects of sex (F_(1,158)_ = 19.60, P < 0.0001) and context (F_(1,158)_ = 73.01, P < 0.0001), however the effect of time, despite marginal, was not significant (F_(4,158)_ = 2.323, P = 0.0590), as were the remaining interactions (all with P > 0.1). Adolescent females display an adequate context discrimination performance in the three first time points, days 2 (**Figure 1.A**) 7 (**Figure 1.B**) and 14 (**Figure 1.C**), then changing to context generalization – not distinguishing between contexts – observed on days 21 (**Figure 1.D**), and 28 (**Figure 1.E**). Surprisingly, and different from adolescent females, these males were simply not able to discriminate the contexts at any time-point, but have learned the fear response, since performance (in % freezing) was comparable to that of females (**Figures 1.F; 1.G; 1.H; 1.I; 1.J**). It is noticeable how the adolescent females maintained memory precision until at least 14 days after training **(Figure 1.K**), whereas males did not show contextual discrimination at any tested time-point (**Figure 1.L**): males actually always displayed a lower discrimination ability compared to the females (**Figure 1.M**). Please, check Table 1 for additional comparisons.

**Table 1.** Statistical analysis to emphasize the specific effects verified in this work: Tukey’s HSD Ps and Cohen’s d values comparing training vs. novel contexts (experiments 1 and 2) and vehicle vs. muscimol (experiment 3) % freezing. Factorial ANOVA for each experiment show some interactions and main effects (see text), which are detailed here (and in the text). (*) P<0.05; *italic font:* strong trend toward significance.

|  | N | P | d | Effect |
| --- | --- | --- | --- | --- |
| Experiment 1 |  |  |  |  |
| 1A | 6,7 | 0.0170 * | 2.1700 | discriminate context A from B |
| 1B | 7,7 | 0.0003 * | 3.4750 |  |
| 1C | 8,8 | 0.0068 * | 2.5350 |  |
| 1D | 8,10 | 0.2447 | - | contexts indistinguishable |
| 1E | 9,10 | 0.6166 | - |  |
| 1F | 8,9 | 0.9950 | - |  |
| 1G | 13,14 | 0.1394 | - |  |
| 1H | 13,13 | 0.9999 | - |  |
| 1I | 6,6 | 0.9999 | - |  |
| 1J | 8,8 | 0.9881 | - |  |
| Experiment 2 | training vs novel |  |  |  |
| females | 7,7 | 0.0074 * | 2.5359 | discriminate context A from B |
| males | 12,12 | 0.9975 | - | contexts indistinguishable |
| ovx females | 10,10 | 0.4855 | - |  |
| Experiment 3 |  |  |  |  |
| females vehicle vs females muscimol | 10,10 | 0.0560 * | 1.7920 | females treated with muscimol differ from all groups |
| females vehicle vs males muscimol | 7,10 | 0.0129 * | 0.3533 |  |
| females muscimol vs males muscimol | 10,7 | 0.0411 * | 1.3349 |  |

### 3.2. Retrieval of a precise memory in female adolescent rats depends on sex hormone levels 14 days after training: ovariectomy accelerates memory generalization to a performance similar to males

In this set of experiments, we checked for the involvement of sex hormones in the maintenance of memory precision in adolescent females. To this end, ovariectomized adolescent females of the same age were tested 14 days after training, in order to compare their performance to that of males and intact females also at 14 days after training – at the suggested transition point for females. Two-way ANOVA indicated group-context interaction (F_(2,52)_ = 3.477, P = 0.0383) and main effect of context (F_(1,52)_ = 13.23, P = 0.0006). In **Figure 2** we see that, while intact (non-ovariectomized) adolescent females can clearly discriminate between the contexts, ovariectomized (OVX) subjects seem to anticipate fear memory generalization, since there is no significatively different % freezing between the training and the novel contexts, resembling what was observed in males. Please, check Table 1 for additional comparisons.

### 3.3. Retrieval of a precise memory in adolescent female rats depends on hippocampus integrity 14 days after training: muscimol infused into the dorsal CA1 region blocks retrieval of fear memory in females, but has no effect upon males

The transformation hypothesis predicts that contextual discrimination memories are precise soon after training since hippocampus is readily available to mediate retrieval, while a remote retrieval tends to become less discriminatory as consequence of their permanent storage in the cortex and progressive independence from the hippocampus (Winocur et al., 2007; Winocur & Moscovitch, 2011). To verify whether memory retrieval was hippocampus-dependent, CA1 area of the dorsal hippocampus was inhibited by the infusion of GABA_A_ agonist muscimol infused prior to testing (in the training context). The specific time-point chosen to investigate memory retrieval was also 14 days after training, when despite already generalized in adolescent males, it still retains some precision in females (Figure 1). Our prediction was that memory retrieval would not be affected by hippocampus inactivation in males, whereas in females it should be impaired.

Two-way ANOVA found sex-treatment interaction (F_(2,30)_ = 4.227, P = 0.0486; females, N=10,10, males, N=7,7) and a main effect of sex (F_(1,30)_ = 6.765, P = 0.0143), but no effect of treatment (F_(1,30)_ = 1.872, P = 0.1814), confirming our expectations **(Figure 3)**. The interaction means different sexes display very different effects, in this case, the visibly amnestic effect caused by muscimol in the female adolescents. Please, check Table 1 for additional comparisons.

## DISCUSSION

Our findings show that, despite being able to adequately retain an aversive experience over different training-test intervals (**Figures 1.F-J** and **1.L**), adolescent males were surprisingly unable to discriminate between similar, yet different contexts. It is as if the aversive and the spatial components of memory were processed at different paces, separately, suggesting a late maturation of this hippocampal cognitive function compared to females of the same age (Giedd et al., 1997; Hu et al., 2013; Lynch et al., 2019). Performance of adolescent males in the training context on day 2 revolved around 50-60% of freezing time (the same as females), and a similarly high recall was observed in every other interval, but males also expressed the same level of fear in the novel context, the very same context females recognize as “different” from the training one in recent memory tests (2 to 14 day intervals). Since in our experimental setup males never display context discrimination, it is not possible to know whether they actually had the ability and did not express it, or they simply lack it to begin with: anyway, we assumed the behavior of “expressing the same level of fear in two different contexts” as an instance of memory generalization as well. Notice that the DI analysis aimed to highlight the lower discrimination ability of adolescent males compared to females of the same age (**Figure 1.M**). The fact that adolescent males can remember (aversion) but do discriminate (contexts) well is intriguing, but can make sense if we speculate about a putative adaptive role, at this age, as a compensatory skill in the face of the characteristic risky behavior frequently exhibited by male adolescent mammals (Ellis et al., 2012).

Adolescent females, on the other hand (**Figures 1.A-E** and **1.K**), exhibited a precise discriminatory memory at least up to 14 days after training, undergoing generalization afterwards. Thus, while there was a clear transition from discriminatory to generalized memory between post-training days 14 and 21 in adolescent female rats, nothing similar took place for their male counterparts.

In order to investigate the involvement of sex hormones in the behavioral change observed in female adolescents compared to male subjects of the same age, a group of ovariectomized females was trained and tested after 14 days later: whereas intact females were able to discriminate between contexts with precision, ovariectomized subjects did not as if context generalization emerged earlier (at a level similar to that of adolescent males) (**Figure 2**), and point to a possible role of female hormones in memory quality, as proposed by others (Keiser et al., 2017; J. Lynch et al., 2013a; J. F. Lynch, Winiecki, et al., 2016). Notice that the controls for OVX females were intact ones, not submitted to sham surgery. We kept it this way based on our controls, in which there was no difference in freezing levels between OVX and non-OVX (intact) females in the training context of experiment 2 (see **Figure 2**) – and even compared to females from experiment 1 (**Figure 1.C**). We thus believe that neither the surgical procedure, nor anesthesia interfere with the effectivity of our memory retention measure, % freezing, and should not come as a surprise considering that those invasive procedures took place at least 14 days before the experiments began.

The last set of experiments showed that the memory precision displayed by adolescent females depends on the integrity of the CA1 region of the dorsal hippocampus, since its bilateral blocking with pretest muscimol was able to impair memory retrieval in females – but not in males, 14 days after training **(Figure 3)**. These results support the idea that a precise memory, in order to be retrieved, needs the hippocampus, exactly what was expected if a reliable systems consolidation process were taking place (Rudy, 2009; Wiltgen et al., 2010a).

Detailed memories are known to depend upon the hippocampus (de Oliveira Alvares et al., 2012; Wiltgen et al., 2010), while generalized ones seem to rely on cortical networks. The standard model of systems consolidation predicts that, with time, the engram becomes progressively independent of the hippocampus and ends up permanently stored in neocortical areas such as the anterior cingulate cortex (Dudai & Morris, 2013; Martin et al., 2005; Jorge A. Quillfeldt et al., 1996; Jorge Alberto Quillfeldt, 2019; Squire et al., 2004, 2015; Squire & Alvarez, 1995). In previous works, we have shown that memory in adult male rats simultaneously exhibit (a) a precise context discrimination and (b) hippocampus dependency up to 28-35 days after training, in agreement with other studies in the literature and consistent with the idea of two concomitant, interdependent subprocesses behind systems consolidation (Crestani et al., 2018; de Oliveira Alvares et al., 2012; Haubrich et al., 2016; Wiltgen et al., 2010). This is why we think adolescent females underwent *full* systems consolidation, while it is senseless even to suppose that systems consolidation was taking place in same age male rats, due to their enigmatic “always-generalized” response.

Here we used the very same training and novel context employed in previous papers studying adult male rats (de Oliveira Alvares et al., 2012; Haubrich et al., 2016), in order to ensure they were easily distinguishable in recent memory tests. With those same conditioning boxes, we have shown that female adolescents display precise memory, i.e., they discriminate the two contexts at least up to 14 days after training, while adolescent males were unable to distinguish them at any of the studied training-test intervals (**Figures 1.A-J**). We interpret the memory failure of the males as a qualitative deficit only – precision is impaired – since there is no actual amnesia: in fact, both male and female animals displayed, at all times, a good recall of the training context, the place where the aversive learning took place (see **Figures 1.K** and **1.L**). Despite the good retention, male subjects reacted with similar fear levels both to the training and the novel context, which we considered to be a generalized response. However, it is advisable to avoid adjectives when talking about generalization, since this particular cognitive phenomenon can equally be the result from loss, or from gain of function. Compared to precision (as in context discrimination), generalization can be taken to be a degraded version of a formerly better trace representation. Nevertheless, from an adaptive point of view, it might as well be a way to facilitate learning via transfer of already acquired information in order to survive in a complex environment (Shepard, 1987).

If this difference in memory capacity represents a developmental sexual dimorphism, one possibility is that the two groups of animals have distinct rates of hippocampal circuitry maturation, something also suggested elsewhere (see, e.g. (Giedd et al., 1997; Hu et al., 2013; K. M. Lynch et al., 2019). Since this difference involves the spatial component, notice that rat spatial navigation systems develop between P20 and P45, being fully functional only at the end of this time window (Scott et al., 2011; Tan et al., 2017): it might well be the case that males and females in late adolescence actually mature their place cells networks at different paces, which opens an interesting avenue for future studies.

The absence of effect of muscimol in males might be simply due to something like the GABAergic connections not being fully established yet **(Figure 3)**. The investigation of this aspect goes much beyond the scope of the present study, however our data at least suggest a role of sex hormones in this process, since ovariectomized females end up responding earlier in a generalized way. Previous studies demonstrated that gonadectomy prior to puberty can alter the organization of neural circuits (Koss et al., 2015) and modify behavioral expression (Delevich et al., 2019; Kercmar et al., 2014). This might also be the case for males, in which testosterone levels are known to increase only after day P40, reaching adult levels at around P75 : since our subjects were trained between P42-P49 – close to the beginning of that developmental window – they might not have yet reached the optimum adult male hormone levels during the synaptic consolidation phase of this contextual fear memory: this also might explain why a less detailed memory was formed. This avenue, however, was not fully explored here, and surely demands further investigation.

Acute and chronic administration of estrogen was shown to accelerate fear memory generalization in adult female rats (J. Lynch et al., 2013a; J. F. Lynch, Winiecki, et al., 2016), while in adult males the exact opposite takes place – testosterone favors memory precision (J. F. Lynch, Vanderhoof, et al., 2016). For our adolescent subjects, just as it seems to be the case for males, our females may have not yet reached the typical adult levels of circulating estradiol at the age of their training and/or test (Vetter-O’Hagen & Spear, 2012), but their hippocampal circuitry is probably more mature than that of males at that time-point, since they are already able to form a precise, hippocampal-dependent trace. Thus, the reduction of estrogen levels by the OVX (**Figure 3**) might be the main cause of this apparent acceleration of generalization, quite the opposite of what caused the generalized responses in adult females for the abovementioned authors (J. Lynch et al., 2013b). In sum, adolescent females can have the benefit of more precise context discrimination memory due to the still reduced estrogen levels (El-Gendy et al., 2019), but then, with time-dependent systems consolidation taking over, comes generalization.

This is very different from what was observed in adult females by others. Even disconsidering some methodological differences between our findings and theirs, the comparison might be useful, since the observed effects appear in blunt opposition: while for adult female rats estradiol replacement (of OVX subjects) leads to generalization, gonadalectomized females (with reduced estradiol) show precise discrimination; in our case, intact adolescents display precise discrimination whereas OVX subjects generalize contexts. We do not know the stage of development of the CA1 circuitry at the training moment in any of the sexes, thus it would be premature to speculate in more detail. Also, with the different training-test intervals it might be interesting to separate the discussion of hormone levels acting upon the training session (acquisition and consolidation) from the possible effects upon the test session, since they might differ a lot, implying different intrinsic mechanisms due to developmental changes.

Finally, in complex studies searching for sex differences, one can never be too careful in considering the relevant variables (Maney, 2016), and context fear conditioning is admittedly a simple behavioral model, despite being adequate to bring phenomena such as systems consolidation to light (Sutherland et al., 2010). One limitation is the use only of one behavioral index – % freezing – when much more is expected to be going on. This might be compensated by measuring additional variables, such as, for instance, darting behavior, a common fear response to footshocks found in female rats that would compete for freezing time (Gruene et al., 2015). However, we did not notice any saliency of this behavior, and since most of our female subjects displayed a good retention of the aversive experience, it was not quantified here.

All things considered, the fact remains that female adolescent rats (a) exhibit the full set of confirming evidence of a *bona fide* systems consolidation process – the change from memory precision to generalization coinciding with the progressive independence from the hippocampus, and (b) this switch can be anticipated by ovariectomy. On the other hand, (c) adolescent male rats, despite being able to learn the task, were not able to distinguish between contexts, always exhibiting a generalized fear response (so that it is not possible to demonstrate the occurrence of systems consolidation). Actually, (d) these male subjects display, at all times, a lower discriminatory capacity compared to females of the same age, suggesting interesting developmental differences in terms of hippocampal circuitry maturation in different sexes. To our notice, this is the first report in the literature of memory systems consolidation in adolescent female rats - and one that can also be accelerated by the reduction of sex hormones through ovariectomy. It is also unprecedented that adolescent male rats, despite remembering the aversive experience well, were not able to discriminate contexts with any precision.

## Author Contributions

APC – Conceptualization, Formal analysis, Investigation, Project administration, Writing – original draft, Writing – review & editing, Visualization, Methodology

FNL – Conceptualization, Investigation, Writing – review & editing, Methodology

MAC – Investigation, Writing – review & editing, Methodology

BP – Investigation, Writing – review & editing, Methodology

KTKG – Investigation, Writing – review & editing

LOA – Conceptualization, Funding acquisition, Resources, Writing – review & editing, Methodology

JAQ – Conceptualization, Funding acquisition, Project administration, Resources, Formal analysis, Supervision, Writing – original draft, Writing – review & editing, Methodology

## Funding acknowledgement

These experiments were supported by CNPq/Brazil, Capes/Brazil and FAPERGS/Brazil fellowships and grants.

## Acknowledgments

We thank Mrs. Zelma Regina Vasconcelos de Almeida for her always resourceful and kind technical assistance.

## Notes

### Competing Interest Statement

The authors have declared no competing interest.

